# AirSurf-*Lettuce*: an aerial image analysis platform for ultra-scale field phenotyping and precision agriculture using computer vision and deep learning

**DOI:** 10.1101/527184

**Authors:** Alan Bauer, Aaron George Bostrom, Joshua Ball, Christopher Applegate, Tao Cheng, Stephen Laycock, Sergio Moreno Rojas, Jacob Kirwan, Ji Zhou

**Affiliations:** Earlham Institute, Norwich Research Park, Norwich, NR4 7UZ, UK; Plant Phenomics Research Center, China-UK Plant Phenomics Research Centre, Nanjing Agricultural University, Nanjing 210095, Jiangsu, China; School of Computing Sciences, University of East Anglia, Norwich Research Park, Norwich, NR4 7TJ; National Engineering and Technology Center for Information Agriculture, MARA Key Laboratory for Crop System Analysis and Decision Making, Jiangsu Key Laboratory for Information Agriculture, Nanjing Agricultural University, Nanjing 210095, Jiangsu, China; G’s Growers Limited, Ely, Cambridgeshire, CB7 5TZ

**Keywords:** Keywords AirSurf, lettuce, ultra-scale field phenotyping, deep learning, image analysis, precision agriculture

## Abstract

Aerial imagery is regularly used by farmers and growers to monitor crops during the growing season. To extract meaningful phenotypic information from large-scale aerial images collected regularly from the field, high-throughput analytic solutions are required, which not only produce high-quality measures of key crop traits, but also support agricultural practitioners to make reliable management decisions of their crops. Here, we report AirSurf-*Lettuce*, an automated and open-source aerial image analysis platform that combines modern computer vision, up-to-date machine learning, and modular software engineering to measure yield-related phenotypes of millions of lettuces across the field. Utilising ultra-large normalized difference vegetation index (NDVI) images acquired by fixed-wing light aircrafts together with a deep-learning classifier trained with over 100,000 labelled lettuce signals, the platform is capable of scoring and categorising iceberg lettuces with high accuracy (>98%). Furthermore, novel analysis functions have been developed to map lettuce size distribution in the field, based on which global positioning system (GPS) tagged harvest regions can be derived to enable growers and farmers’ precise harvest strategies and marketability estimates before the harvest.

## Introduction

As an important source of vitamins, minerals, and trace elements, leaf vegetables play crucial roles in human nutrition^1^. Lettuce (*Lactuca sativa* L., a plant of the Asteraceae family), one of the most common staple vegetable foods, has a wide range of tastes, textures, and shapes that satisfy diverse customer needs^2,3^. Recent research indicates that lettuce consumption has positive effects on the reduction of cardiovascular disease and chronic conditions, because it provides nutrients such as vitamin A, *Beta-carotene*, folate, and iron content to support human growth and health^4,5^. While lettuce is an important and nutritional crop, fluctuating environments can increase the fragility of its production^6^. For example, the bad weather in Spain in early 2017 led to retail prices for lettuce products to nearly triple in UK supermarkets^7^. Severe weather not only can cause supply shortage, but also affects the crop quality. According to studies on lettuce growth and development^8–10^, at newly planted phase (i.e. from cotyledons unfolded to three true leaves stage), young crops require cool and damp weather to develop into high-quality products after transplanting from greenhouse to the fields; whereas lettuce leaves can rapidly become bitter and inedible if the plant growth is accelerated by ambient temperature at the head maturity phase (i.e. before flowering). Because of the dynamic nature of lettuce production, the actual yield of lettuces in commercial operations is around 70-80% of the planted quantity^11,12^. To ensure consistency of supply and quality, it is important for growers and farmers to closely monitor lettuce growth and development, so that prompt and reliable agricultural practices can be arranged under today’s fluctuating agricultural conditions^13^.

Commercially, lettuce production offers an attractive economic profitability in comparison to many other Agri-Food businesses^14,15^. To date, lettuce-related businesses are worth billions of dollars and employ hundreds of thousands of permanent and seasonal workers globally. According to the Food and Agriculture Organisation of the United Nations^16^, European vegetable growers alone produced 2.95 million tonnes of lettuce (and chicory) in 2016, a total annual value of €2.5 billion. Spain, the largest lettuce producer in Europe, is exporting approximately €420 million worth of lettuce products every year; Germany, France and the UK are the three largest markets for lettuce consumption in Europe, with a combined import of €350 million annually^17^. To serve diverse consumer tastes as well as improve the actual yield of lettuces, lettuce breeders are constantly introducing new varieties to the market, from the dense head (i.e. iceberg type) to the notched or frilly leaf varieties^18,19^.

Further down the fresh produce supply chain, the planning and efficiency of many essential Agri-Food activities are largely dependent on the maturity date and marketability of different sizes and quality of crops^20^. Logistics, trading, and marketing need to be organised several weeks before the harvest^21^; moreover, the booking and reservation of lettuce distribution, agricultural equipment, and commercial plans with retails are often determined between H1 (soft and spongy head) and H3 (mature and compact head) stages. So, crop can be harvested at the right time, with maximised yield^22,23^. To reliably measure and estimate potential yield (e.g. the number of lettuce heads) and associated crop quality (e.g. lettuce size categories) for better marketing and supply chain management, growers and farmers are continuously seeking new technologies to assist them with better and more precise crop management decisions^24,25^.

As a relative newcomer to life sciences, machine learning (ML) related techniques use statistics and sparse representations to progressively build computational procedures to accomplish specific tasks such as data classification, feature selection, clustering, and predictive modelling^26,27^. Although the unfamiliarity with ML often prevents plant researchers from effectively employing ML and its related technology in biological studeis^28–30^, many cases indicate that ML is the key to success in addressing a variety of data-driven challenges in life sciences, if appropriately labelled training data^31^, suitable learning algorithms^32^, and well-defined missions can be arranged. Some of the cases are: (1) the analysis of big genomic data for annotation, assembly, and gene regulatory networks^31^, (2) the classification of DNA and protein sequences for genetic and genomic studies^33^, and (3) the prediction of genome and phenome patterns based on high-dimensional feature datasets^34^.

In this article, we present a cross-disciplinary approach that develops a new analytic software tool to perform automated ultra-scale field phenotyping of iceberg lettuce. Our research and development (R&D) activities integrates ultra-scale normalized difference vegetation index (NDVI) aerial imagery, modern computer vision, state-of-the-art deep learning (i.e. convolutional neural networks, CNNs), supervised machine learning, modular software engineering, and commercial lettuce production into an open-source image analysis platform called AirSurf-*Lettuce* (AirSurf-L). The platform is capable of performing phenotypic analysis of millions of lettuces across the field. A CNN model trained with over 100,000 labelled lettuce signals has been embedded in AirSurf-L to quantify lettuce heads and their plantation layout using ultra-large NDVI images collected by a fixed-wing light aircraft. Unsupervised ML algorithms were used to classify lettuce heads into three size categories (i.e. small, medium and large). To connect analysis results with marketability and crop management decisions, a novel function has been developed to connect global positioning system (GPS) information with the lettuce size distribution map, based on which a GPS-tagged harvest map was produced to enable efficient and precise harvesting strategies for growers and farmers to increase marketable yield. The analysis results generated by AirSurf-L show a strong correlation between machine counting and specialist scoring. We are therefore confident that our work is promising in assisting vegetable growers and farmers with their precision agriculture management activities. Together with recent advances in unmanned aerial vehicles (UAV) technologies, ground-based remote sensors, and ML-based modelling, AirSurf-L could have great significance to improve the fresh vegetable crop production, distributing and logistics activities before the harvest. Furthermore, with addtional training data, necessary testing and validation, we believe that the analysis platform can be expanded relatively easily to incorporate other crop species such as wheat and rice for ultra-scale aerial crop phenotyping.

## Results

### NDVI aerial imaging and data acquisition

The ultra-large aerial NDVI imagery was acquired routinely (i.e. four-five times per season) using a fixed-wing light aircraft operated by G’s Growers, the second largest vegetable grower in the UK. The flying route and the imaging protocol were designed to facilitate cross-site crop layout assessment, yield prediction (based on vegetation indices), and disease monitoring (Fig. 1A), which has been described previously^35^. In this study, we used a series of collected ultra-large NDVI images (1.5-2GB per image) at 3cm ground sample distance (GSD) spatial resolution, collecting iceberg lettuce signals between H1 and H2 stages (i.e. moderate compact and crushable head), before lettuce leaves were largely overlapped. Experimental fields were located in Cambridgeshire UK, ranging from 10 to 20 hectares, with between 800,000 and 1.6 million lettuce heads in a single field. A field planted with around 1 million lettuce heads (coloured light blue in Fig. 1B) was used in the following sections to explain the analysis workflow of AirSurf-L. A high-level manual yield counting was conducted by the grower’s field specialists during the harvest, which was used to verify and improve the AirSurf-L platform. Also, lettuces in subsections randomly selected from experimental fields were scored manually by laboratory technicians, which were also used as training data for the CNN model.

**Figure 1:**
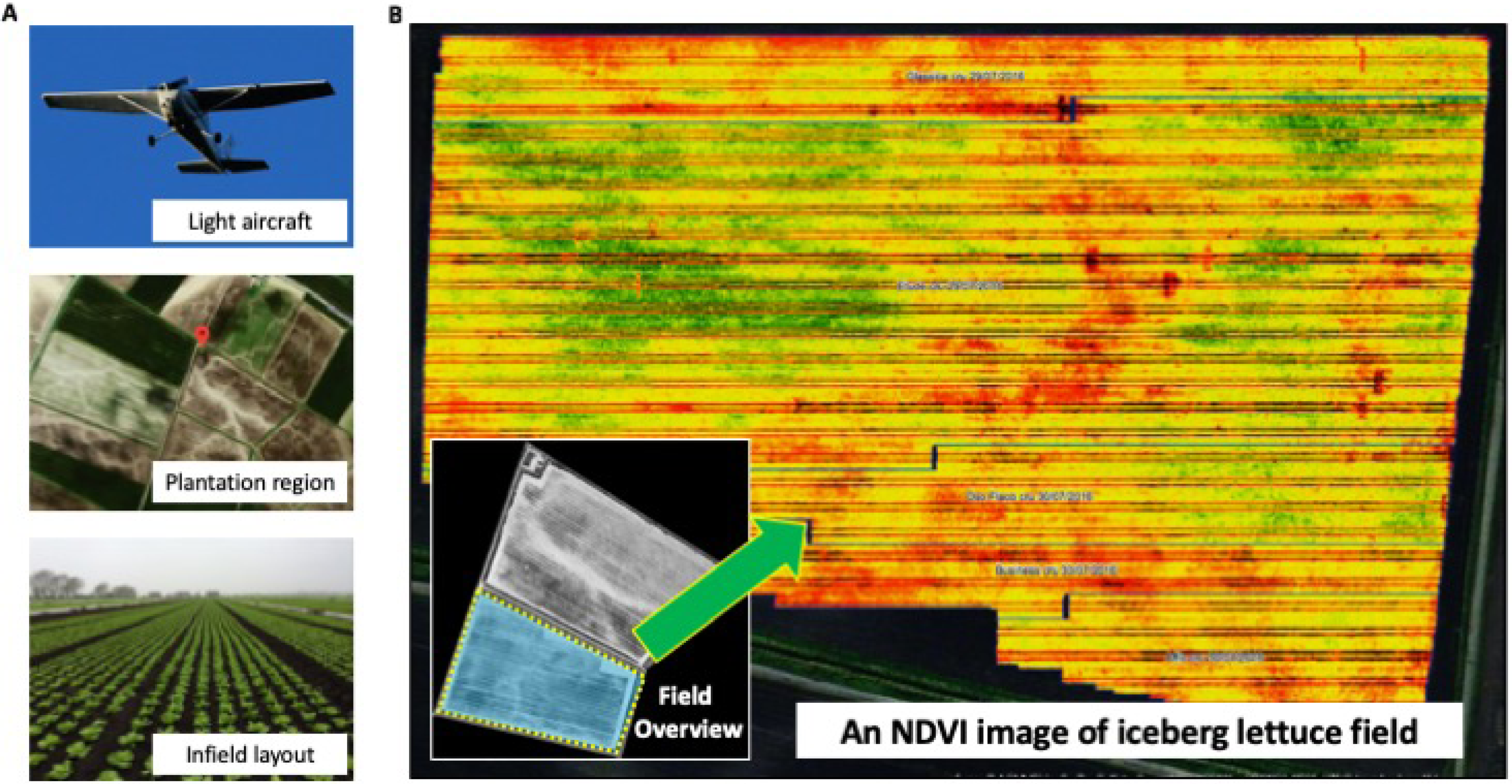
Ultra-large NDVI aerial imaging accomplished routinely through a fixed-wing light aircraft operated by G’s Growers. **(A)** The flying route and aerial imaging were designed to facilitate cross-site crop layout assessment and yield prediction. **(B)** A series of ultra-large NDVI images at 3cm GSD spatial resolution were acquired to record 0.8-1.6 million lettuce heads per field, at H1 and H2 stages.

### The analysis workflow of AirSurf-L

The analysis of yield-related phenotypes was based on NDVI signals of iceberg lettuces across the field. Figure 2 shows a high-level analysis workflow of AirSurf-L, which consists of five steps: data input, image calibration and pre-processing, ML-based traits analyses, results visualisation, and quantifications of yield-related phenotypes. *Step 1* accepts raw NDVI images as gray-level imagery datasets. As pixels with extremely high NDVI signals usually have overflowed intensity values (i.e. black pixels in Fig. 2A), a pre-processing step (*Step 2*) is designed to calibrate raw NDVI images, so that intensity distribution can be normalised and overflowing pixels can be corrected. In this step, an algorithm called contrast limited adaptive histogram equalization (CLAHE)^36^ is applied to increase the contrast between the foreground (i.e. lettuces) and background (i.e. soils) in a given NDVI image (Fig. 2B). Additional File 1 provides pseudo code and explanations of this step.

**Figure 2:**
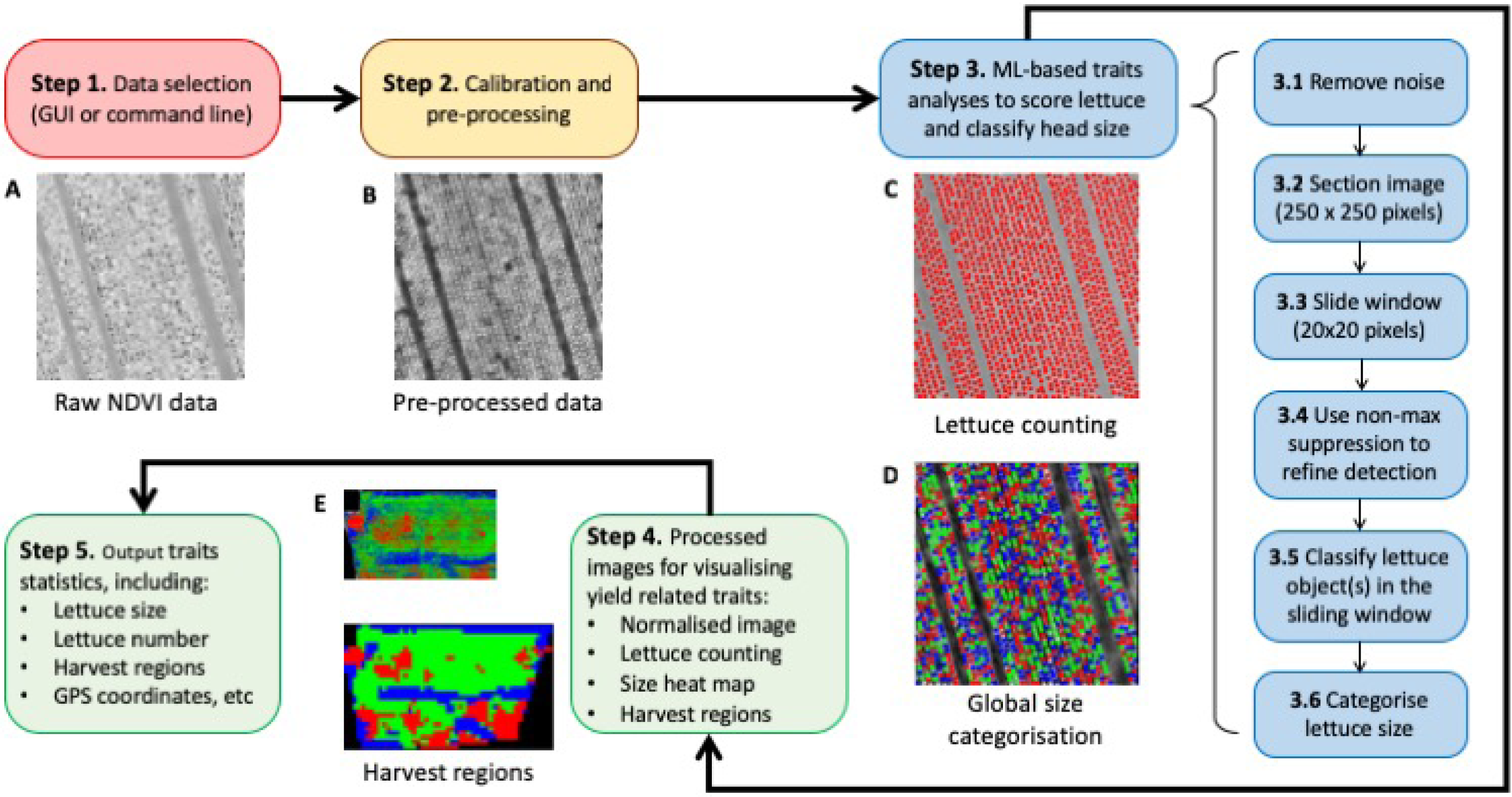
A high-level analysis workflow of AirSurf-Lettuce. **(A)** Step 1 accepts raw NDVI images as input imagery data (pixels with extremely high NDVI signals are overflowed). **(B)** Step 2 pre-processes the raw NDVI images to calibrate intensity distribution and correct overflowing pixels. **(C&D)** Step 3 carries out ML-based traits analyses to quantify lettuce number and classify head size across a given NDVI image. (E) Steps 4&5 visualise and export statistics of the traits analyses detection, including yield-related phenotypes such as lettuce counting, size distribution, and harvest regions, and associated GPS coordinates.

*Step 3* carries out ML-based traits analyses that quantify lettuce number (Fig. 2C) as well as classify head size (Fig. 2D). It includes six steps: removing noise signals, partitioning a given image into sections (250 x 250 pixels) for local analysis, producing a sliding window (20 x 20 pixels) to traverse within a sectioned image, using non-max suppression to detect lettuces, and classifying recognised lettuces into three size categorises (i.e. small, medium and large). The analysis result is visualised in *Step 4*, where lettuce counting, size distribution map, and GPS-tagged harvest regions are saved as processed images (Fig. 2E). At the final step (*Step 5*), statistics of yield-related traits are saved in a comma-separated values (CSV) file, including lettuce counts per field, lettuce size distribution, lettuce number and size measures within GPS-based grids, harvest regions, and their associated GPS coordinates (Additional File 2). To enable users to carry out the above analysis workflow, a GUI-based software application has been developed (Supplementary Fig. 1).

### Data construction for model training and testing

To generate a sound training and testing dataset for ML-based image analysis, we randomly selected dozens of patches of a given field and manually labelled each lettuce in the patch with a red dot (Supplementary Fig. 2). Then, each dot is identified by a bounding box, which is then used to build the learning model. A training dataset with over 100,000 20x20-pixel labelled images has been created, amongst which 50% are lettuces and the remaining are background signals such as soil, edges of the field, and other non-lettuce objects. Following a standard CNN segmentation approach^37^, we designed a sliding window function to go through a given image to divide foreground and background signals, splitting lettuce and non-lettuce objects. Training and testing datasets are equally balanced. Validation sets are used alongside training sets to verify the performance of the model as well as to prevent overfitting in model training, which is also an important step to allow us to fine-tune hyperparameters of different learning layers^38^.

### Neural network architecture

Similar to AlexNet^39^, a CNN-based learning architecture was established using the labelled datasets. Figure 3A demonstrates the architecture of the CNN model, including (1) a convolutional (Conv2D) layer with 32 filters and a 3x3 kernel, with a rectified linear unit (ReLU) as the activation function, and batch normalisation to accelerate the learning process to enable higher learning rates^40^; (2) the same block is then repeated together with a max pooling layer to down-sample input using a 2x2 kernel based on the assumption that useful input features could be contained in sub-regions; (3) after that, a second convolutional block is constructed, consisting of a Conv2D layer with 64 filters, a 3x3 kernel, a ReLU activation, and batch normalisation; (4) finally, this block is repeated, followed by another max pooling layer (with a 2x2 kernel) to complete the learning procedure. After the convolutional layers, layers are connected to a fully connected layer of size 512, which is followed by a dropout layer with a 50% chance. To complete the learning architecture, a binary output generates the probability of whether a given bounding box (20x20 pixels) contains a lettuce signal. If the probability equals or is close to 1.0 (i.e. 100%), it indicates that it is highly likely that the bounding box contains a complete lettuce head (Fig. 3B). The above architecture is commonly applied to vision-based object detection problems^41^. The training and validation accuracy and loss curves are reported in Figure 3C, showing that the model converges in only 10 epochs. To avoid overfitting, the stopping criterion was designed to ensure that the validation accuracy is higher than the training accuracy. By doing this, we can ensure the generalisation of the model. When training the CNN model, the labelled data was also divided equally into train and validation sets.

**Figure 3:**
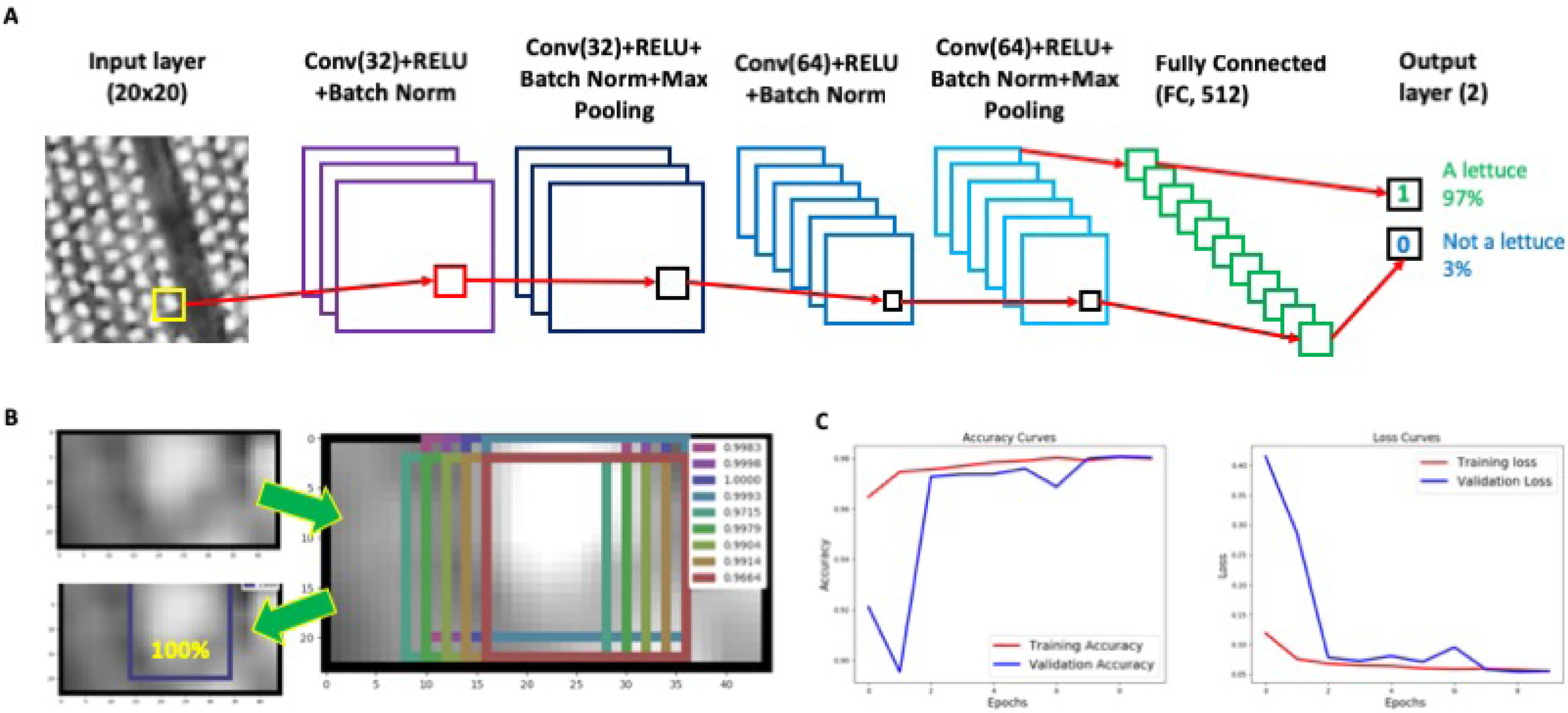
A CNN-based learning architecture established for lettuce counting. **(A)** The architecture of the trained CNN model, which generates a binary output representing the probability of whether a yellow bounding box contains a lettuce signal. **(B)** If the probability is close to 1.0, it indicates that it is highly likely that the bounding box encloses a lettuce. **(C)** The training and validation accuracy and loss curves of the model.

### Phenotypic analysis of lettuce heads

After a CNN classifier was trained, we used it to recognise and classify lettuce signals in ultra-large NDVI images. The six-step approach discussed before (Fig. 2) is followed. However, during the testing and implementation, we found that the CNN classifier could wrongly score lettuces as a lettuce head is extremely tiny in an orthomosaic image (e.g. 11,330x6,600 pixels for a 7-hectare field when GSD is 3cm, which can contain over half million lettuce heads). To resolve this issue, we have designed a two-step approach: (1) sectioning the whole image into many 250x250 pixels sub-images, and (2) using a fix-sized bounding-box (20x20 pixels) as a sliding window (with a stepping parameter of 5 pixels to reduce the sliding calculation) to prune the detected lettuce objects in each sub-image.

Another reason that caused the CNN classifier’s wrong detection is overlapped lettuce objects. Even in a sub-image, overlapped lettuces could be detected repeatedly by the classifier. Hence, we employed a non-maximum suppression (NMS) algorithm^42^ to rectify the detection result. NMS uses probabilities to order the detected lettuce objects in a sub-image. After the sliding window function is performed and many small patches have been identified in a sub-image, the NMS algorithm computes an overlap coefficient to determine how to retain these patches. As lettuces are relatively well-spaced, bounding boxes surrounding a complete lettuce signal will be retained, whereas partially covered signals will be removed. To select the best overlap parameter for the NMS, a gradient descent method is formulated. Additional File 3 explains the NMS algorithm and its implementation in AirSurf-L.

### Results improvement and size categorisation

Initially, the training data selected was chosen randomly across the field. Using the data, AirSurf-L can capture a broad range of sizes and orientations of lettuces with varying intensities. However, when applying the initial CNN model, it failed to recognise lettuces in very bright regions and overly count lettuces in very dark regions (e.g. approximately 50,000 lettuces were wrongly detected in the one-million-head field, Fig. 4A). To resolve this issue, we enhanced the training datasets by manually labelling an additional 500 lettuce signals within very bright or very dark regions. Then, newly labelled data was inserted into the training datasets to retrain the model through the online-learning approach^43^. The improved model (available on our GitHub repository, see Availability of supporting data) was tested on different experimental fields again and has dramatically enhanced the detection result (Fig. 4B).

**Figure 4:**
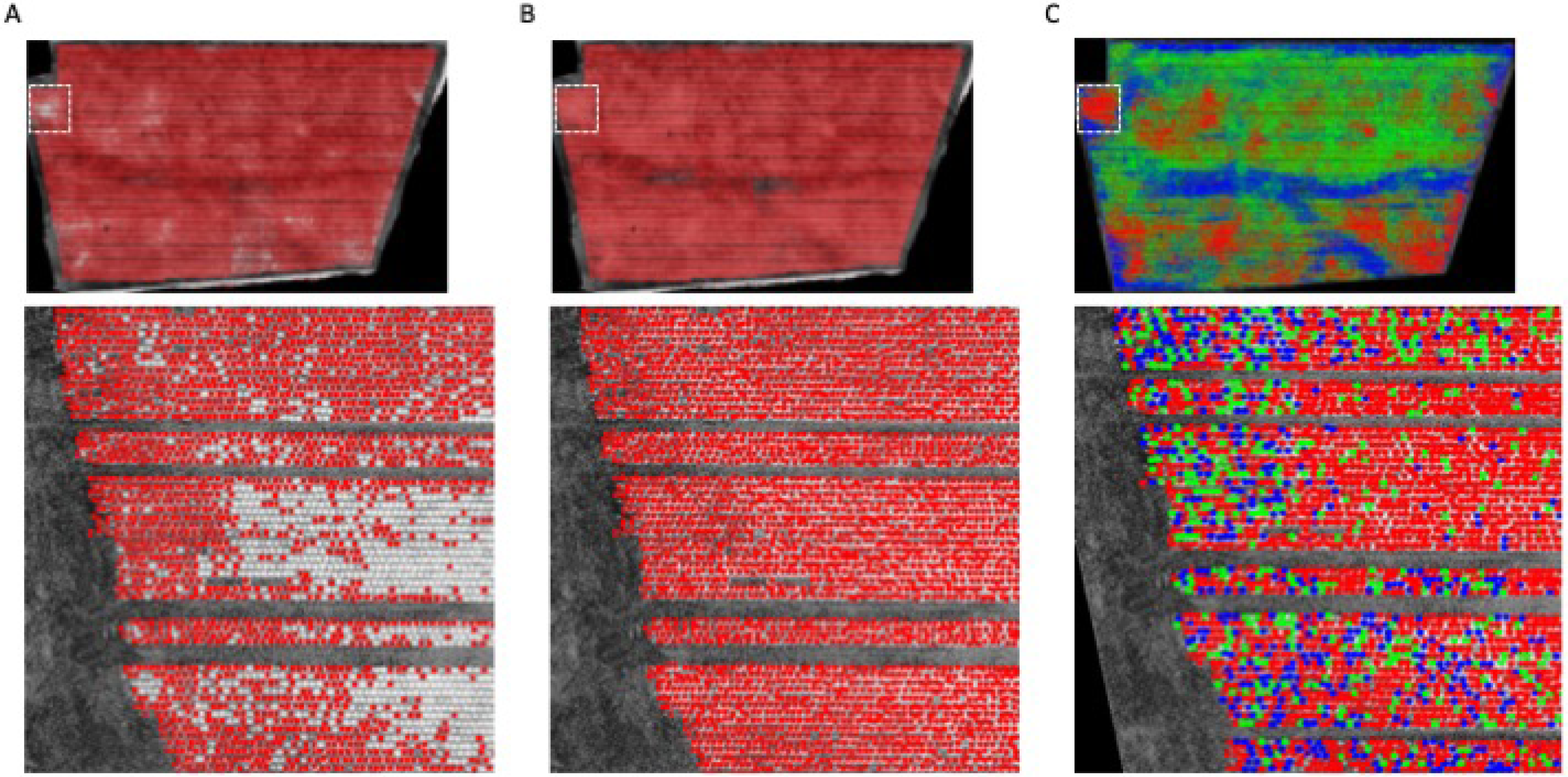
The improved results of the CNN model and the size classification of lettuce heads. **(A)** Wrongly detected lettuces in very bright regions and overly counted lettuces in very dark regions, in a one-million-head field. **(B)** Enhanced training datasets to retrain the model using the online-learning approach, which led to much better detection results. **(C)** A predefined colour code (small is coloured blue, medium is coloured green, and large is coloured red) is assigned to each recognised lettuce head across the field.

Identified lettuces are individually analysed to determine their associated size category. The size classification is based on intensity and contrast values enclosed by the 20x20 bounding boxes, which is computed using the dot product of the histogram of pixel intensities and a weighted vector towards more pixel-based contrast values (see Methods). The assumption of this design is that higher NDVI signals likely correlate with higher vegetation indices^44^, which indicates bigger lettuce heads. The categorisation result of all lettuce heads is clustered into three size groups. Each lettuce is then coloured with a predefined colour code (i.e. small is blue, medium is green, and large is red, see Fig. 4C).

### GPS-tagged harvest regions

The final phase of the phenotypic analysis is to define harvest regions based on colour-coded lettuces. Using the size distribution map (Fig. 5A), the field is firstly segmented into many small grids based on the optimal GPS resolution determined by the altitude of aerial imagery (3cm GSD, in our case) as well as the size of the harvester machinery used by the grower. After dividing the field into thousands of grids (Fig. 5B), GPS coordinates of each grid are recorded and each grid is coloured with the most representative lettuce size category. By combining all coloured grids, a GPS-tagged harvest map is produced, representing harvest regions of the field (Fig. 5C). The harvest map is then used for designing harvesting strategies such as guiding a harvester to collect desired sized lettuces or arranging logistics based on the lettuce number and size counting. To facilitate agricultural practices, a result file (in .csv format, Additional File 2) is generated by AirSurf-L at the end of the analysis, containing information of each harvest region, the associated GPS location, lettuce size, lettuce counts, and the location in the field. To satisfy different needs for dissimilar requirements, the size of GPS-based harvest grids can be modified manually in the AirSurf-L software.

**Figure 5:**
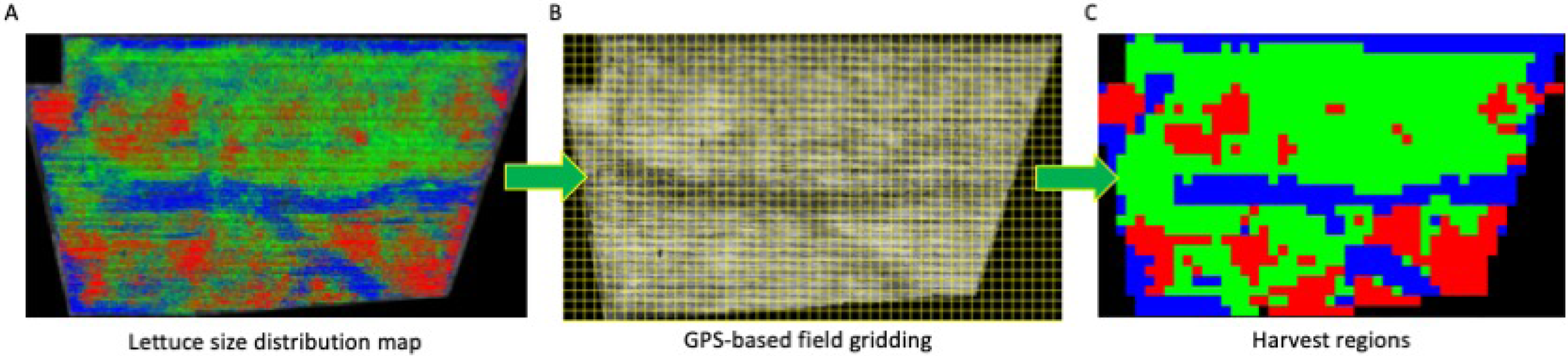
A GPS-based harvest map based on lettuce size classification. **(A)** A colour-coded lettuce size distribution map. **(B)** The field is segmented into thousands of grids based on the optimal GPS resolution and the size of the harvester machinery. **(C)** Grids are coloured with the most representative lettuce size category across the image, representing harvest regions of the whole field.

Figure 6 uses Python-based 3D Matplotlib library^45^ to show the GPS-tagged harvest map. When AirSurf-L reads an NDVI image, it first computes the number of lettuce heads and associated size categories on the image (Fig. 6A). Then, by 3D visualising the relationship of GPS-based field grids, the number of lettuces in the grid, and the representative size category (Fig. 6B), a dynamic 3D bar chart is generated to present lettuce number using the z axis, infield layout (both columns and rows) using both x and y axes, and the representative lettuce size using the predefined colours (Fig. 6C). Through the 3D plot, users can zoom into one sub-region of the field to check detailed yield-related traits within each infield grid and plan harvesting strategies accordingly.

**Figure 6:**
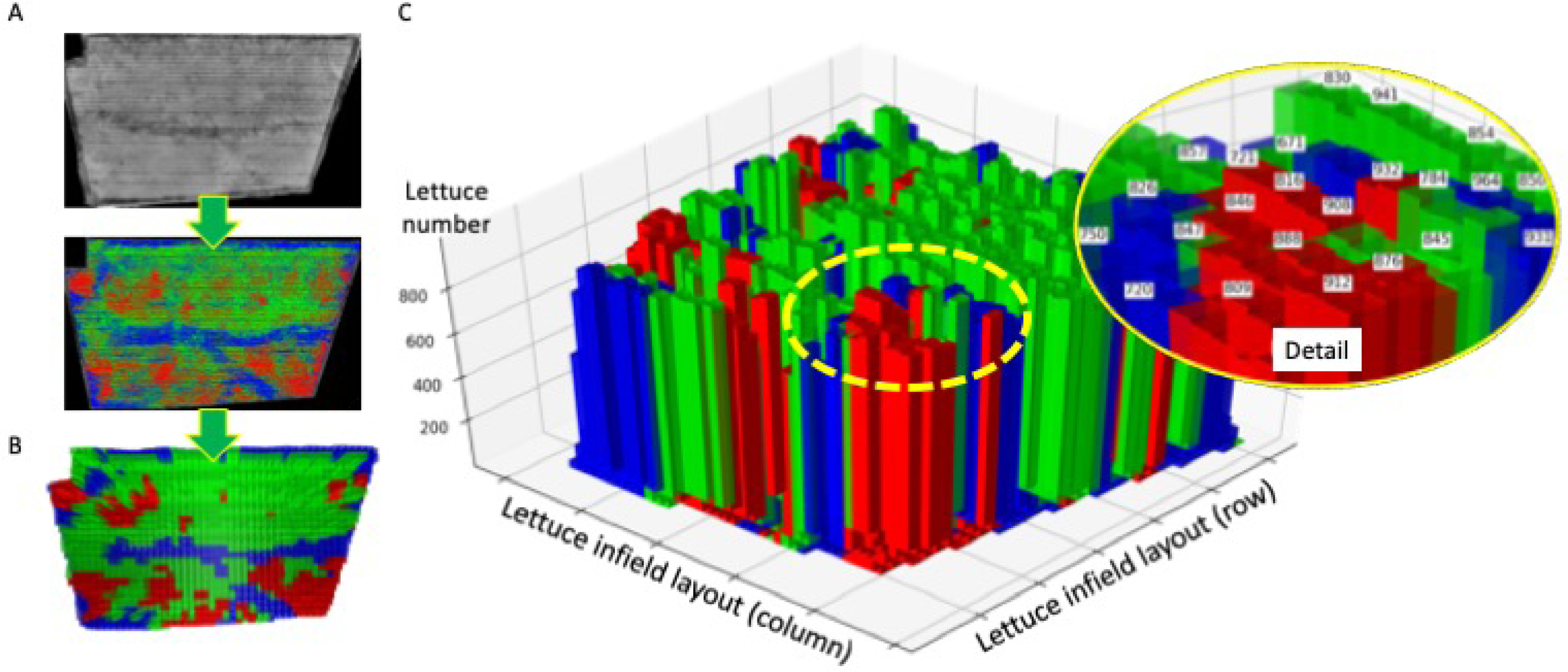
3D visualisation of lettuce harvest regions. **(A)** AirSurf-L reads an NDVI image and exports a lettuce size distribution map. **(B)** 3D visualising GPS-based field grids to represent the number of lettuces, representative size categories. **(C)** A dynamic 3D bar chart is generated to present the relationship between lettuce number, infield layout, and the representative lettuce size.

### Validation of the platform

To verify AirSurf-L and the generalisation of the learning model, we have applied the platform to count and classify lettuce heads in three unseen experimental fields in Cambridgeshire, UK (Figs. 7A-C). These fields contain around 700,000-1,500,000 lettuces and are located in different sites around the county. Traits such as the number of lettuces per field and associated size categorisation quantified by the platform were compared with industrial estimates, showing a highly correlated phenotypic analysis (<5% difference). Besides the field-level comparison, we also randomly selected different sizes of subsections in a given experiment field to evaluate AirSurf-L more precisely. We split these subsections into two sets (i.e. 36 small regions and 21 large regions), where the small regions have less than 400 lettuces and the large ones contain greater than 900 lettuces heads. After that, laboratory technicians manually counted lettuce heads within these regions. The correlation between the manual and AirSurf-L counting shows that, for the small regions, the difference between the human and automatic counting is approximately 2%; for the large regions, the average difference is around 0.8%. Supplementary Figure 3 reports the strong correlations (R^2^ = 0.98) between human and automatic counting in both regions.

**Figure 7:**
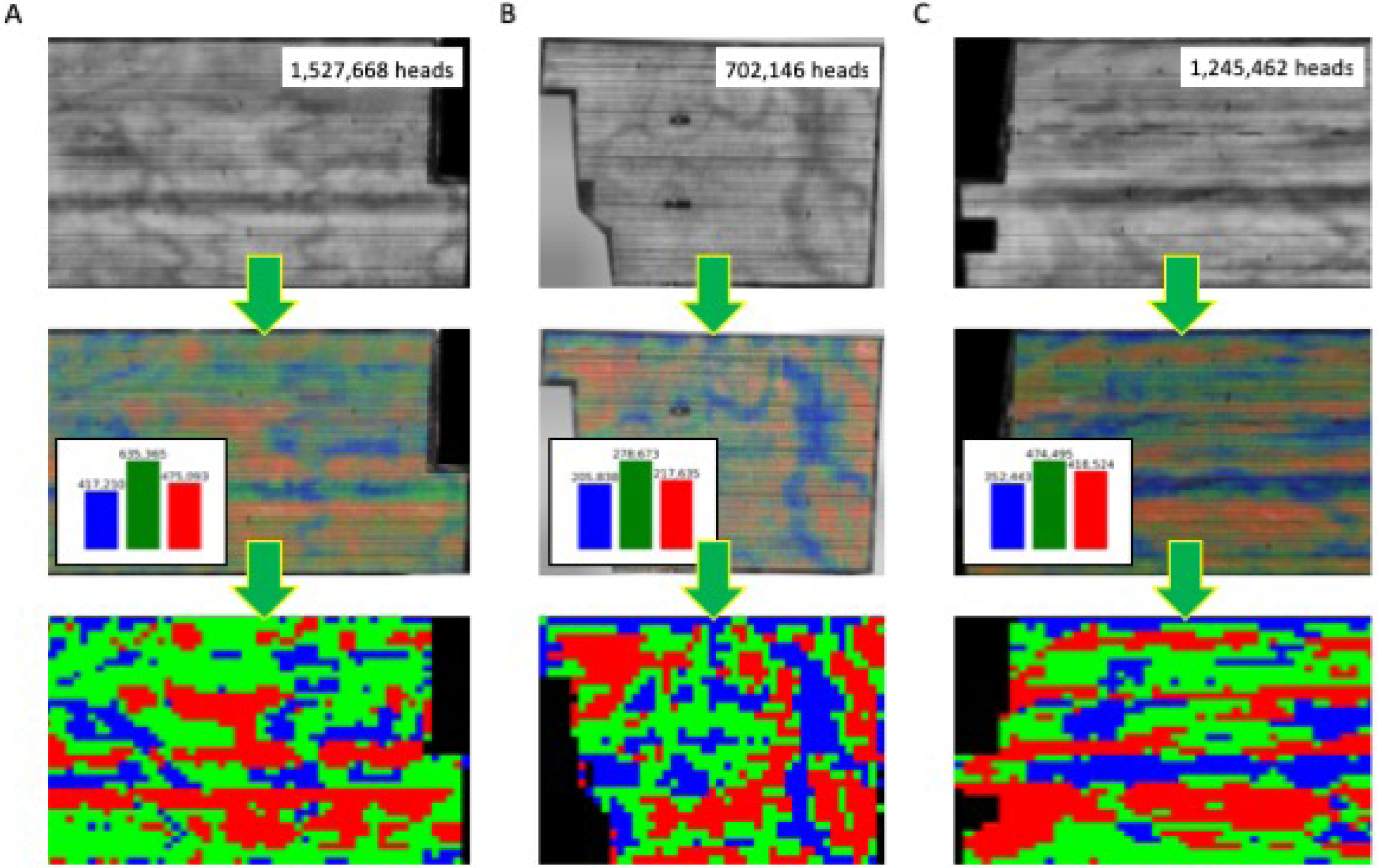
Applying AirSurf-Lettuce to count and classify millions of lettuce heads in three plantation fields across the Cambridgeshire, UK. **(A-C)** AirSurf-Lettuce is applied to count and classify millions of lettuce heads in three plantation fields in the Cambridgeshire, UK.

## Discussion

Traditionally, measuring infield crops on a large scale is very time-consuming and labour-intensive. It often requires destructive techniques, potentially error-prone manual counting, or estimates of traits that are key to yield production or crop quality^46^. Recent advances in machine learning (including deep learning) and computer vision (CV) based techniques have led to an explosion of phenotypic analysis that is rapidly improving our abilities to mine phenotypic information from large and complicated phenotyping datasets^47–49^. New data-driven analytic approaches are also changing plant phenomics research – collecting big data (i.e. phenotyping) is no longer the bottleneck, instead how to extract biologically relevant information (i.e. phenotypic analysis) from big data has become the current challenge^50–52^. Hence, along with the development of aerial imaging and remote sensing technologies, it has become increasingly noticeable that the integration of scalable data collection, high-throughput phenotypic analysis, and predictive modelling are key to crop research and precision agriculture^52–54^.

### A combined research effort

AirSurf-L addresses a specific challenge in ultra-scale field phenotyping and precision agricultural practices through combining aerial NDVI imagery, CV, ML, and moldular software engineering, with commercial lettuce production. The software platform automates the measurement of millions of lettuces in the field and our industrial partner has contributed key ideas of how to connect research-based phenotypic analysis with real-world agriculture problems. As a cross-disciplinary project, our project collaborators came from different backgrounds and hence many efforts were made on understanding the requirements at the project initiation phase. Also, the academia-industry project setting required a more agile R&D progress, because computational technologies and industrial requirements were constantly changing while the project was still ongoing. From the project development, one of the most valuable lessons we learned is that requirements and implementation are unlikely to be clarified at the beginning and more efforts shall be made towards mutual understanding. Similar to the case reported previously^55^, all project parties need to be adaptive with changeable requirements due to the dynamic nature of such a project; additionally, a successful integration of project stakeholders requires all parties to manage expectation, mutual trust, and, more importantly, clear communication channels.

### Machine learning in plant phenomics

Another aim of this work is to further ML-and CV-based software solutions in plant phenomics. High-throughput plant phenotyping is a fast-growing research domain, covering many disciplines, from plant breeding, cultivation, remote sensing, to computing sciences^56,57^. The modulated software development allows us to apply different open-source learning architectures^58^ (e.g. Scikit-Learn and the TensorFlow frameworks) and CV algorithms^59,60^ (e.g. OpenCV and Scikit-Image libraries), when implementing AirSurf-L. Notably, it is worth pointing out that ML is *not* a silver bullet for phenotypic analysis, because: (1) learning algorithms could perform badly if training datasets are not well-labelled; and, (2) although ML performs well in segmentation and classification if target objects are well-defined, there is still a big gap between object recognition and traits analyses. Meaningful phenotypic analysis not only requires sufficient biological understanding to define target traits in a logical manner, but also needs bespoke algorithms to engineer features so traits can be soundly extracted. Hence, biological questions, CV, data analysis, and software engineering shall be considered collectively with ML techniques when resolving plant phenomics problems.

### Limitations of the platform

Besides the high-accurate phenotypic analysis results presented in this article, there are still limitations of the platform need to be considered: (1) AirSurf-L has been tested with top-view iceberg lettuces mainly at H1 and H2 stages, which means that analysis error could increase if there are too many overlaps between lettuce heads, e.g. from H3 stage onwards. (2) As AirSurf-L has only been tested with NDVI imagery, it is therefore important to add new functions to the platform to incorporate other vegetation indices measured through multi-and hyper-spectrum imaging sensors. (3) As precision agriculture management decisions are normally based on imagery, soil and climate conditions, AirSurf’s results will be more reliable, if soil and climate data can be integrated in the analysis. (4) The method was tested and validated in lettuce fields in a number of geographic locations following a standard aerial imaging procedure, data collected from different sites via varied aerial imaging strategies (e.g. different angles, altitudes and GSD) could improve the soundness and compatibility of the platform.

### Prospects for crop research and precision agriculture

The open-source software development opens up the potential for computational biologists to include new learning models and analytic functions for other staple crops such as wheat and rice. For example, the plant density of wheat is closely related to the yield due to its influences on the allocation of water, light and fertilisers; however, it is not feasible to quantify the plant density solely using ground-based RGB imagery^61^. Hence, utilising the ultra-scale NDVI aerial imagery and object recognition methods embedded in AirSurf-L, it is likely possible to quantify wheat plants at the emergence stage in different farming sites, which not only can benefit the assessment of sowing performance, emergence rate, and plant distribution, but also will help breeders and cultivation researchers make early predictions of the grain yield of different wheat genotypes in field experiments. From a precision agriculture perspective, monitoring individual plant such as a lettuce head will enable accurate monitoring of crops during key growth stages across a plantation site. It can provide growers with the real number of crops in the field, based on which yield for harvest availability and management plans can be quantified instead of estimated. The calculation of infield crops will also lead to more accurate agricultural inputs, facilitating automated variable-rate application of fertiliser, weed control, and pesticides through tractor software system with a precise crop distribution map^62^. More interestingly, the close monitoring of key yield-related traits can also be used to guide farmers and growers to reduce variability of agrichemical applications and irrigation in different fields, leading to increased harvest yield and better operating profit margin^63^. Finally, the AirSurf-L approach fits in the cost-effective category in precision agriculture. The platform utilises existing aerial imagery data routinely performed by the growers and farmers, which means that no extra data collection cost is required by this approach and hence the adoptability of the technology, an important factor for new Agri-Tech solutions to be accepted by the Agri-Food sector^53^.

## Conclusions

AirSurf-*Lettuce* automatically measures infield iceberg lettuces using ultra-scale NDVI aerial images, with a focus on yield-related traits such as lettuce number, size categories, field size distribution, and GPS-tagged harvest regions. The analysis results are close to the manual and industrial counting and can be used to improve existing crop measures and estimates. By monitoring millions of lettuces in the field, we demonstrate the significant value of AirSurf-L in ultra-scale field phenotyping, lettuce size distribution mapping, precise harvest strategies, and marketability estimates before harvesting. We believe that our algorithm design, software implementation, lessons learned from applying ML-and CV-based algorithms, and the academic-industrial R&D activities will be highly valuable for future plant phenomics research projects that are destined to be dynamic and cross-disciplinary. Finally, with continuous development work, we are confident that the analytic platform is likely to support the Agri-Food sector with a smarter and more precise crop surveillance approach of vegetable crops and therefore lead to better precision agriculture management decisions.

## Methods

### Ultra-large field NDVI imagery and experimental fields

The NDVI imaging sensor used is an industrial standard camera described previously^35^. The aerial imaging was carried out by a ‘Sky Arrow’ light aircraft, the lightest weight class (Very Light Aircraft, VLA) of any commercial aircraft that is allowed for commercial work. The VLA let the pilot to fly with very little fuel, less than an average farm vehicle while driving around the crops. Using VLA at 1000 feet (around 305 metres) in the sky, vast areas can be covered at a flight speed 180-200 km/hour. The NDVI sensor can gather ultra-large crop imagery datasets at very low operating costs, as the VLA can carry 45 kilograms of payload to cover four or five fields in a single flight. This aerial imaging approach can also be used to understand the spectral changes associated with key disease conditions. The NDVI lettuce signals used in this study were captured at H1 or H2 growth stage (moderately compact and crushable head, when lettuce leaves are not largely overlapping with neighbours). The experimental fields are operated by G’s Growers near Ely UK, ranging from 10 to 20 hectares with at least 0.5 million lettuce heads in a single field. A rough manual yield estimate was produced by specialists during the harvest, which were used for testing and improving AirSurf-L.

### Dataset preparation

To generate a sound dataset for machine learning-based image analysis, we randomly selected 60 patches of the field of varying sizes, each containing between 300 and 1,000 individual lettuce heads, and manually labelled each lettuce in the selected patches. Each labelled lettuce is extracted as a 20x20 pixel image representing a single lettuce head. We then used these images, along with images that did not correspond to a lettuce head, to train a CNN classifier to recognize and separate potential lettuces from the ultra-large field images. The pixels contained within the potential lettuces were used for further phenotypic analysis of lettuce size. To format the manually labelled dataset for building the model, we created another training dataset with over 100,000 20x20 pixel images, among which 50% are lettuces and 50% are background signals. The background images were selected using regions other than the labelled lettuces across the field together with a non-overlapping sliding window function to extract background patches. These images are then split into two equally balanced training and validation sets.

### Construction of deep neural network architecture

We built our deep neural network based on the architecture of AlexNet. We used a shallower architecture as opposed to AlexNet and other modern deeper architectures for several reasons: (1) the size of our dataset is relatively small for deep learning studies, where larger and deeper networks tend to require bigger training datasets; (2) additionally, ours is only a binary classification problem as opposed to the ImageNet classification task; (3) larger neural networks often require more time to train, which can be slower to execute and not feasible to train the model without specialised hardware such as GPUs. In our case, we wanted a relatively simple, but powerful model that could execute in a broad range of environments and in a timely manner.

Like AlexNet, we used rectified linear units (ReLU) as our activation function, which is now a common standard in CNNs. This is because ReLU reduces the vanishing gradient problem^19^. After each convolutional layer, we also perform batch normalisation. This reduces the covariance shift, which helps ensure that the model generalises well and the network converges faster. Finally, we included two max pooling layers to reduce the problem into smaller samples. Other architectures might use more max pooling layers, but our input images were segmented and hence quite small. In order to avoid too much information loss from the training procedure, we trained the CNN on our datasets until the validation accuracy was greater that the training accuracy. The training and validation accuracy and loss are reported in Figure 3, where it is shown that the model converged in only 10 epochs. More importantly, to avoid over fitting, the stopping criterion was set for when the validation accuracy is higher than the train accuracy.

### Size categorisation

After AirSurf-L identifies a list of square pixel patches containing single lettuces, it is important to perform automatic unsupervised size categorisation. Lettuce sizing in this work is split into three size categories: small, medium and large; however, the method can be easily changed to classify more size categories. The pixel regions are extracted from the image and then NDVI values are put into bins with similar pixel values. Originally, the histogram included 10 bins that are evenly spread across the value range, i.e. 0-255. However, treating all pixel values equally performed poorly in practice. We therefore included two important aspects in the size categorisation. Firstly, the lower NDVI surrounding value does not determine the actual size of the lettuce; secondly, the higher NDVI values are much more important in indicating size. As such, we created a geometric pattern of cut-off values for each bin. These were: 64, 128, 160, 192, 208, 224, 232, 240, 244, 248, 250, 252, 253, and 254. With these cut-off values, most of the background pixels were captured in the first two bins, with increasing granularity as the values approached the maximum of 255.

Having transformed the pixel regions into a series of histogram count vectors, we were able to compare regions and cluster the patches into groups. The count vectors are grouped into three distinct sizes by using k-means clustering with k set to three. They are sorted into size order through calculating the dot product between the weight vector and the cluster centres count vector. These sorted values then determine which cluster corresponds to which size, and subsequently, applies to each lettuce. Three colours are used to indicate size categories: blue for small, green for medium, and red for large.

### Common Pitfalls

The CNN trained on a set of approximately 100,000 images. Despite the reported training and validation accuracies being quite high, in practice the network performed poorly because it could not distinguish lettuces in patches where most lettuces appear particularly bright. As the initial training datasets were chosen randomly, not enough representative samples from extreme regions were selected during the training. Without sufficient training data, the network was undercounting by 5% in large fields. To solve this problem, we manually labelled further 500 lettuces and added them to the training dataset. The neural network was retrained and converged faster than the previous iteration. The algorithm was updated with the new model with improved results. The above training issue could be a common pitfall for many deep-learning analytic solutions, because key features were constructed by learning algorithms instead of engineered. Many learning models were vulnerable when facing up to totally undefined datasets.

## Availability and requirements

Project name: AirSurf-Lettuce with G’s Growers

Project home page: https://github.com/Crop-Phenomics-Group/Airsurf-Lettuce

Operating system(s): platform independent

Programming language: Python 3

Requirements: Packaged for both Mac and Windows

License: BSD-3-Clause available at https://opensource.org/licenses/BSD-3-Clause

## Abbreviations

Comma-separated values: CSV
computer vision: CV
convolutional neural networks: CNNs
deep learning: DL
global positioning system: GPS
ground sample distance: GSD
machine learning: ML
non-maximum suppression algorithm: NMS
normalized difference vegetation index: NDVI
rectified linear units: ReLU
the United Kingdom: UK
Unmanned Aerial Vehicles: UAVs

## Availability of supporting data

The datasets supporting the results presented here is available at https://github.com/Crop-Phenomics-Group/Airsurf-Lettuce/releases. Source code and other supporting data are also openly available in the GitHub repository.

## Author contributions

J.Z., A.B., A.G.B. and J.K. wrote the manuscript, S.M.R. preformed the NDVI imaging. J.K. provided harvest information and biological expertise. J.Z., A.G.B., and C.A. designed the analysis algorithms. A.B., A.G.B., C.A. and J.Z. developed and implemented the core algorithms. A.B. and A.G.B. built the deep learning models for AirSurf-L. J.B. packaged the GUI executables. J.Z., A.B., A.G.B., C.A., S.L. and J.B. tested the software. J.Z., A.G.B and A.B. performed the data analysis. All authors read and approved the final manuscript.

## Acknowledgements

The authors would like to thank all members of the Zhou laboratory at EI and Nanjing Agricultural University for fruitful discussions and cross-country collaborations. We thank researchers at John Innes Centre and UEA for constructive suggestions. We gratefully acknowledge the support of NVIDIA Corporation with the award of the Quadro GPU used for this research.

---

**Supplementary Figure 1:**
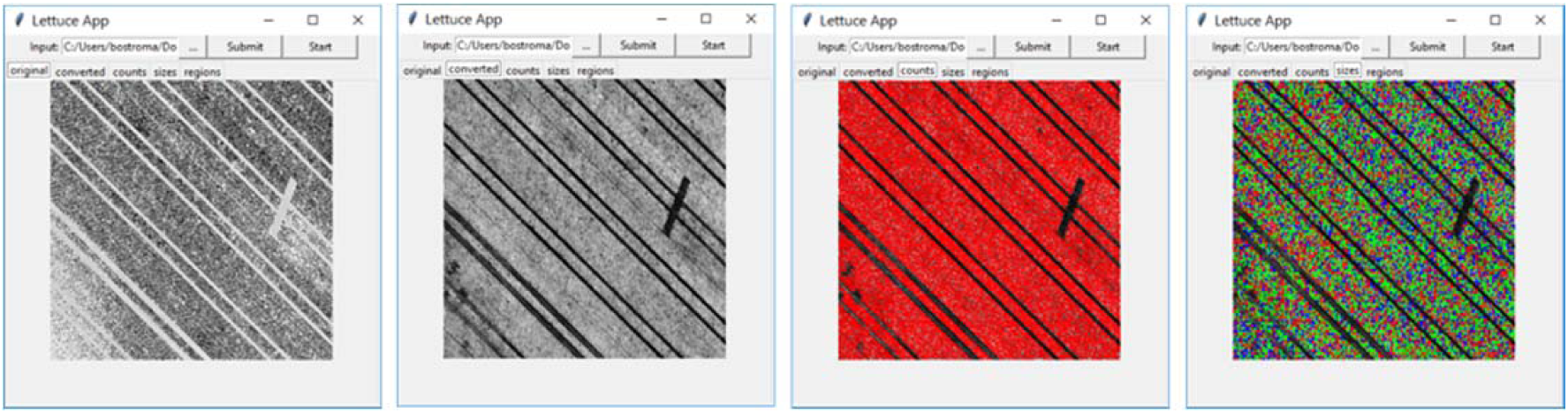
The GUI interface of AirSurf-Lettuce and the analysis workflow.

**Supplementary Figure 2:**
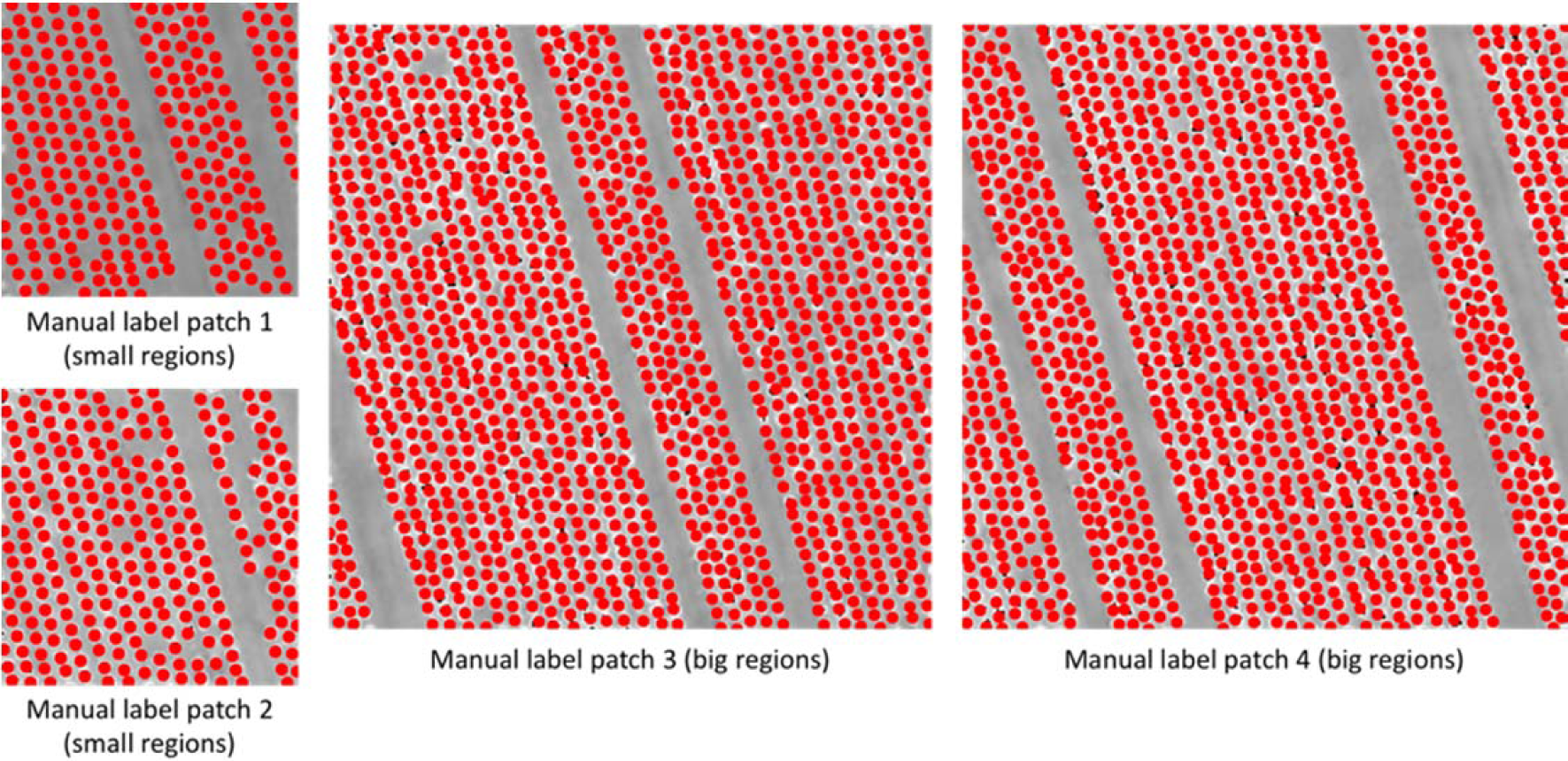
Manually labelled lettuces in randomly selected patches using red dots.

**Supplementary Figure 3:**
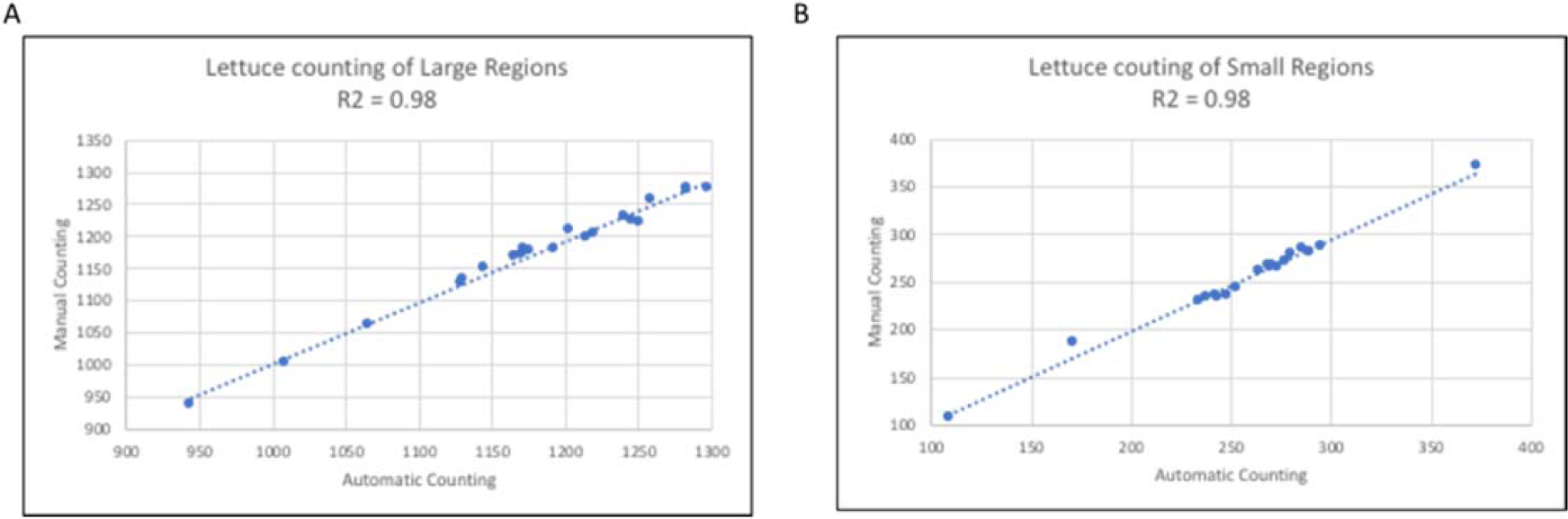
The correlation between human counting and AirSurf-L scoring (R^2^ = 0.98).

